# Tracing Back the Temporal Change of SARS-CoV-2 with Genomic Signatures

**DOI:** 10.1101/2020.04.24.057380

**Authors:** Sourav Biswas, Suparna Saha, Sanghamitra Bandyopadhyay, Malay Bhattacharyya

## Abstract

The coronavirus disease (COVID-19) outbreak starting from China at the end of 2019 and its subsequent spread in many countries have given rise to thousands of coronavirus samples being collected and sequenced till date. To trace back the initial temporal change of SARS-CoV-2, the coronavirus implicated in COVID-19, we study the limited genomic sequences that were available within the first couple of months of its spread. These samples were collected under varying circumstances and highlight wide variations in their genomic compositions. In this paper, we explore whether these variations characterize the initial temporal change of SARS-CoV-2 sequences. We observe that *n*-mer distributions in the SARS-CoV-2 samples, which were collected at an earlier period of time, predict its collection timeline with approximately 78% accuracy. However, such a distinctive pattern disappears with the inclusion of samples collected at a later time. We further observe that isolation sources (e.g., oronasopharynx, saliva, feces, etc.) could not be predicted by the *n*-mer patterns in these sequences. Finally, the phylogenetic and protein-alignment analyses highlight interesting associations between SARS-CoV-2 and other coronaviruses.

## 1 Introduction

Coronaviruses are a large family of RNA viruses that cause diseases in mammals and birds [1]. Coronaviruses have been identified broadly in avian hosts along with various mammals, which include camels, masked palm civets, bats, mice, dogs, cats, and human. The coronaviruses cause respiratory and neurological diseases in hosts [2]. In human, it causes several illnesses, ranging from mild common cold to much severe diseases such as SARS-CoV (Severe Acute Respiratory Syndrome) and MERS-Cov (Middle East Respiratory Syndrome). They are enveloped viruses with a positive-sense single-stranded RNA genome. The genome size of coronaviruses ranges between 26-32 kb [3], the largest among known RNA viruses [4].

The recent coronavirus outbreak in China [5], originating from Wuhan, has been characterized by the identification of a novel type of betacoronavirus SARS-CoV-2 that infects human [6]. In this paper, we restrict our study to those samples collected and sequenced between December, 2019 and January, 2020. In China, 11,791 cases were identified as infected with SARS-CoV-2, and 17,988 were suspected in 34 provinces as of January 31, 2020 [7]. Due to the quick spread of SARS-CoV-2, more than 200 countries and territories around the world have already been marked with the viral infection [8]. It is therefore interesting to trace back whether the origin and spread of this virus has any influence on the change of its nucleotide sequence.

The sequencing of viral genomes has opened up new promises through bioinformatics analyses [9, 10]. With the recent efforts of sequencing the SARS-CoV-2 samples collected across the world, hundreds of genomic samples are becoming available day by day [11]. This has lead to the discovery of new pathophysiological facts about coronaviruses. It is equally interesting to explore the mutations happening in the samples emerging across the world. In this paper, we aim to verify whether the changes of sequence in the samples can reveal the temporal footprints (precisely the month of collection) of this virus.

## 2 Related Work

Since the outbreak of 2019 novel coronavirus disease (COVID-19) in late December, 2019, several studies have already been carried out in understanding the virus and its pathophysiology. The SARS-CoV-2 has been identified by the close genomic observations of clinical samples from patients with viral pneumonia in Wuhan, China [12]. Identification of the molecular mechanism of SARS-CoV-2 is essential to prevent and control the disease. Through next generation sequencing, homology has been observed in the structure of the receptor-binding domain between SARS-CoV and SARS-CoV-2. The SARS-CoV-2 is found to largely deviate from both MERS-CoV and SARS-CoV and is closely related to two bat-derived SARS-like coronaviruses [12].

Another recent work has performed an entropy-based analysis with the diverse genomes and proteomes of SARS-CoV-2 [13]. The results of this study highlighted low varying genomic sequences within the SARS-CoV-2 sequenced samples where the higher variability was observed in at least two nucleotide positions in protein-coding regions. The interplay between the receptor-binding domain (RBD) of spike glycoprotein of SARS-CoV-2 and angiotensin-converting enzyme II (ACE2) complex in human had been extensively studied in several other papers. As the free energy of RBD-ACE2 binding for SARS-CoV-2 is remarkably lower than that for SARS-CoV, the SARS-CoV-2 is epidemically stronger than SARS-CoV [14].

The presence of high prevalence, ecological distribution, tremendous genetic diversity, and frequent alterations of coronavirus genomes have already been implicated in the recurrence of this virus in humans [15]. However, there is hardly any study that links the sequence patterns (precisely *n*-mer distributions here) to the timeline of sample collection. There are many recent resources on coronaviruses due to their repeated outbreak. CoVDBd is a repository of annotated coronaviruses^1^ [16]. There are recent attempts to understand the genome composition and divergence of SARS-CoV-2 [17], however, such studies are restricted to China. Our analysis is more global from this perspective.

## 3 Materials and Methods

The materials and methods used in this paper are all available either from the developers or from us for public use. Additional details are given below.

### 3.1 Dataset Details

We collected the SARS-CoV-2 genomic sequences from NCBI Virus portal^2^ that cover multiple countries. Only those countries that have at least 5 samples were considered and complete genomic sequences were taken. Total 36 sequences (17 samples from China, 5 samples from Japan, and 14 samples from USA) were obtained from the 87 genomic sequences of SARS-CoV-2 available in NCBI. These samples are gathered from multiple patients. For all these genomic sequences, we collected the geographical locations, sources of isolation, and months of collection. The collection month of SARS-CoV-2 samples confirm the presence of the virus in the patients of China in December, 2019 and January, 2020, of USA in January, 2020, and of Japan in January, 2020. The exact collection dates are available for the samples of China (except one) and USA but not for Japan (see Supplementary Dataset 3). The sequences from Japan have recently been removed from NCBI but are still accessible^3^. An inquiry to the submitter regarding the reason of this removal has resulted in no reply so far. The sequences of other types of coronaviruses were collected from NCBI for phylogenetic analysis.

### 3.2 Methods

The Sieve plot was created for geographical location versus isolate (source of viral sample collection) using the Orange toolbox^4^ [18]. The area of each rectangle in this is proportional to the expected frequency of different isolation sources with respect to the countries, while the observed frequency is shown by the number of squares in each rectangle. The Sieve diagram highlights frequencies in a two-way contingency table and compare them to expected frequencies under the assumption of independence. Several classification algorithms were performed that are standard [19]. The classification results were taken from Orange. For this, the default values of the parameters set in Orange were considered. The performance of the classification models are determined in terms of the five evaluation criteria, namely 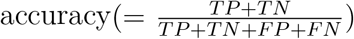, 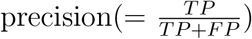, 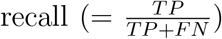, 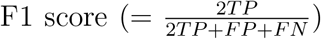, and AUC (area under ROC curve), where *TP, TN, FP* and *FN* stands for the number of true positives, true negatives, false positives and false negatives, respectively.

FASTX (version 36.3.8h Aug, 2019) was used for multiple sequence alignment of DNA sequence of the reference genome of SARS-CoV-2 with the proteins in UnitProt [20]. The parameters for FASTX were: BL50 matrix (15:−5), open/ext: −10/-2, shift: −20, ktup: 2, E-join: 0.5 (0.0832), E-opt: 0.1 (0.0198) and width: 16. The fast family and domain predictions were also obtained using FASTX.

The phylogenmetic tree was built using the tool embedded in NCBI Virus and further processed in the form of a consensus network using SplitsTree (version 5.0.0_alpha). The original input consisted of 269 taxa. The Consensus Network method [21] was used (with default options) so as to obtain 535 splits as compatible.

## 4 Results

The different analyses performed on the SARS-CoV-2 genomic sequences are reported hereunder.

### 4.1 Extraction of Genomic Signatures

The frequency counts for several *n*-mers (for *n* = 1, 2 and 3) were extracted yielding a total 84 (4+16 + 64) features. The frequency values were normalized by dividing with the length of the sequence. The feature similarity across the different samples showed higher importance of shorter *n*-mers (Fig. 1).

**Figure 1:**
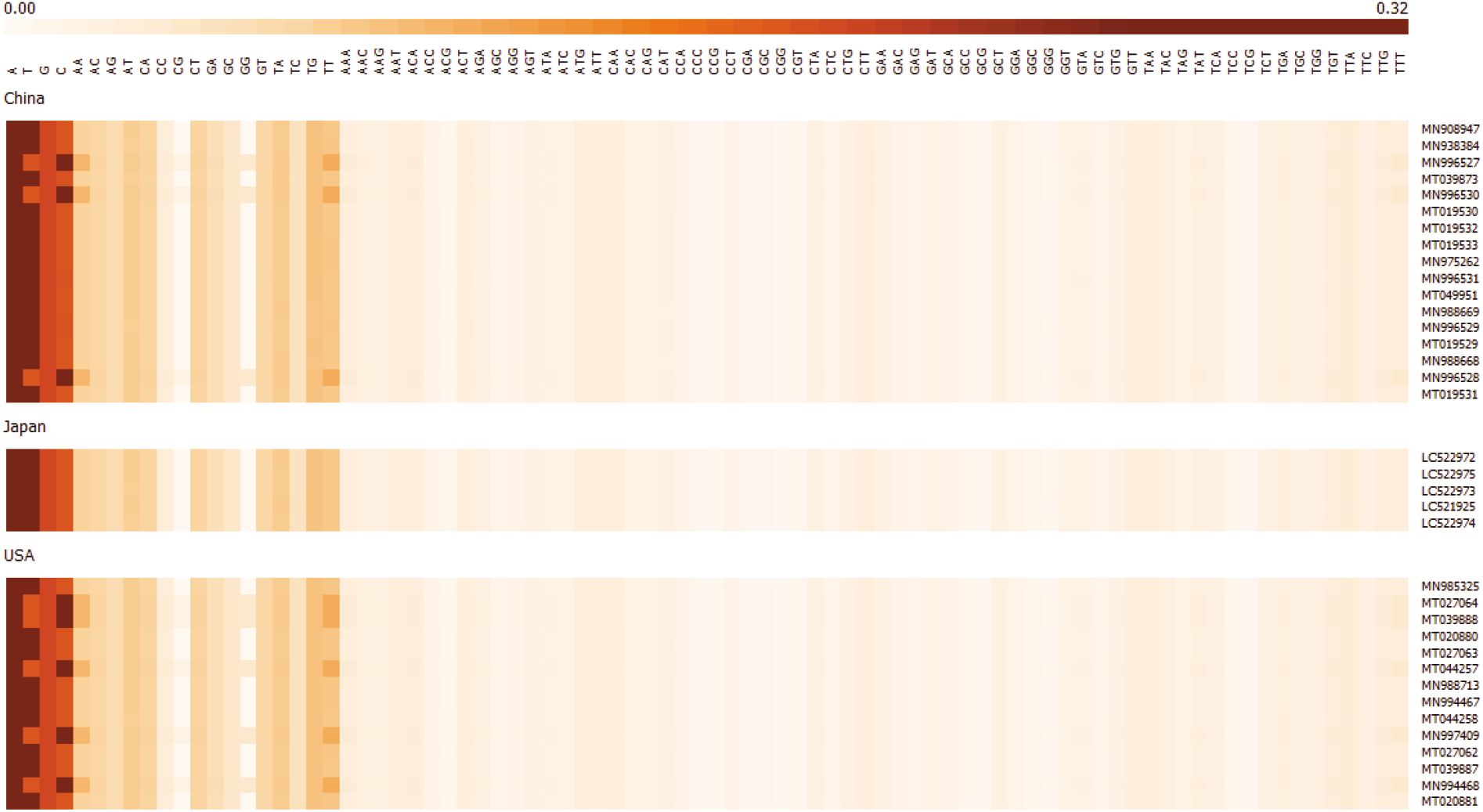
The heatmap reflecting the similarity of *n*-mer distribution across multiple SARS-CoV-2 samples. The samples are grouped by the geographical location from where they were collected.

To test the power of *n*-mer features in distinguishing different geographical locations from where the samples were collected, hierarchical clustering with complete linkage was performed (with number of clusters set to be 3) on the feature data. On labeling the clustering result (Fig. 2), a strong group between the samples collected from Japan was found. The samples from USA and China were highly overlapping. It is interesting to note that all the samples from Japan were collected in late January, whereas those from USA and China were collected much earlier. This highlights a possibility of having a temporal change of SARS-CoV-2 sequences at an early stage.

**Figure 2:**
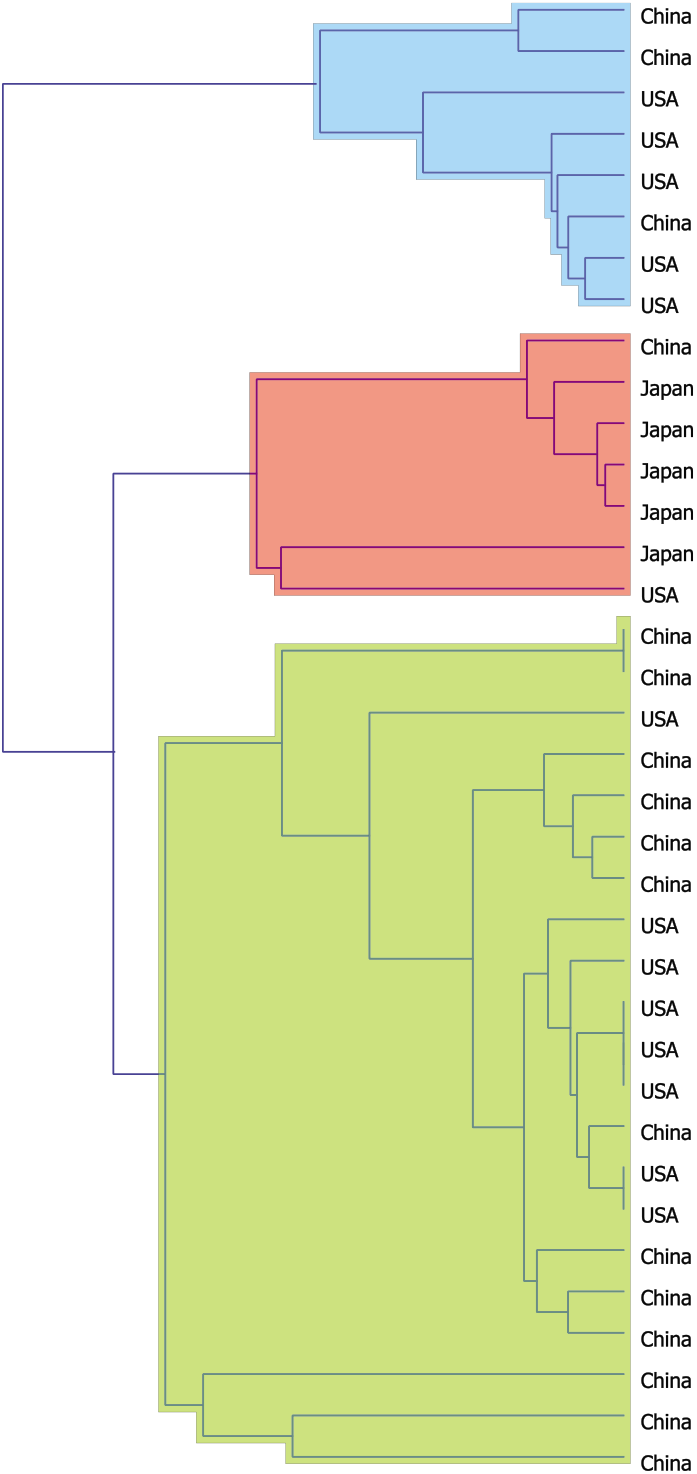
The clusters obtained from the samples based on *n*-mer features by applying hierarchical clustering with complete linkage. The samples from Japan that were collected relatively later came under a single group. Note that, all the samples from Japan have a distinct collection month of January, 2020.

### 4.2 Tracing Back the Temporal Change

As the samples we considered were collected over a period of couple of months, classification results were obtained by setting collection months as the labels against the samples. The collection months December, 2019 and January, 2020 were used as the labels of the samples and multiple classification algorithms were applied (with 5-fold cross validation). The classification results (with 5-fold cross-validation), in terms of classification accuracy, precision, recall, F1 score, and area under the ROC curve (AUC), are reported in Table 1. Close to 78% prediction accuracy was obtained with the Random Forest model, thereby indicating a connection between the changes of *n*-mer in the SARS-CoV-2 sequences over time.

**Table 1:** Performance evaluation of different classification approaches (5-fold cross validation) based on the 84 *n*-mer features to distinguish the months when samples were collected. The best values over a column are shown in bold.

| Classification | Accuracy | Precision | Recall | F1 Score | AUC |
| --- | --- | --- | --- | --- | --- |
| k-NN | **77.8%** | **0.763** | **0.778** | 0.758 | 0.743 |
| AdaBoost | **77.8%** | **0.815** | **0.778** | **0.787** | **0.793** |
| Random Forest | **77.8%** | **0.767** | **0.778** | **0.770** | **0.820** |

As many of the samples from USA and Japan were collected lately, we further studied the importance of both geographical location and temporal signature to distinguish the footprint of SARS-CoV-2. On performing a clustering of samples based on the best features labeled with geographical location and collection time, well segregated groups were found. This highlights a clear distinction of samples based on collection details (Fig. 3). The gradual mutations in SARS-CoV-2 sequences might be the reason of the said distinguishing pattern between the samples collected in December, 2019 and January, 2020. Unfortunately, with the newer (collected in March and April, 2020)) SARS-CoV-2 samples added to the list, this distinctive pattern of the corresponding sequences no more exists. However, it is interesting to note that, our classification attempts consider the class label as month of collection and the data is skewed over the month. In fact, the samples are now getting sequenced with an exponential growth. Therefore, further investigation is required for a deeper understanding of the temporal change of SARS-CoV-2 sequences at a later phase.

**Figure 3:**
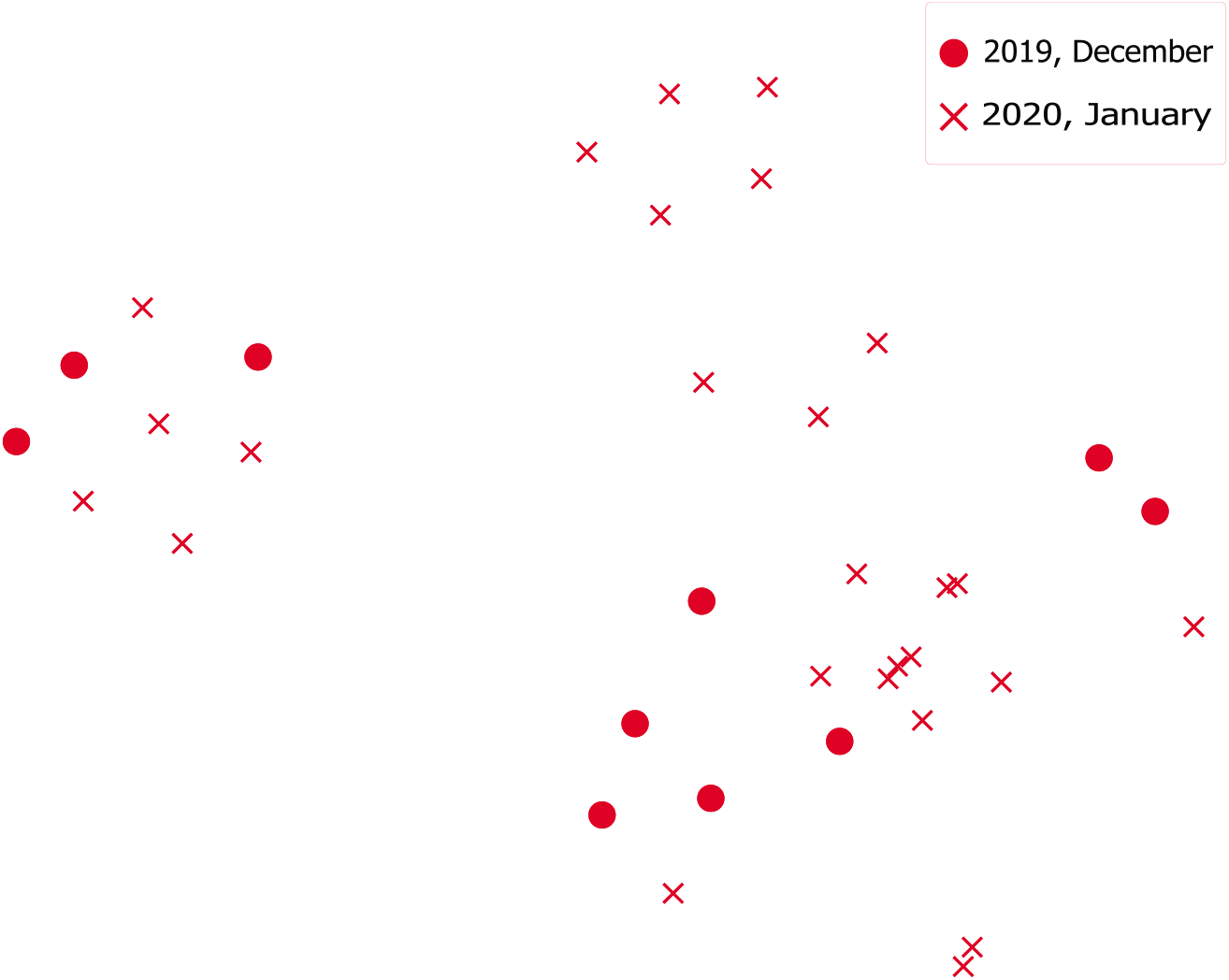
The clustering of samples based on selected features to distinguish collection month.

### 4.3 Relevance of Isolation Sources

Countrywise isolation sources were very much overlapped and do not provide any significant information. The distribution of samples across the different isolation sources as collected from each geographical location are shown in Fig. 4.

**Figure 4:**
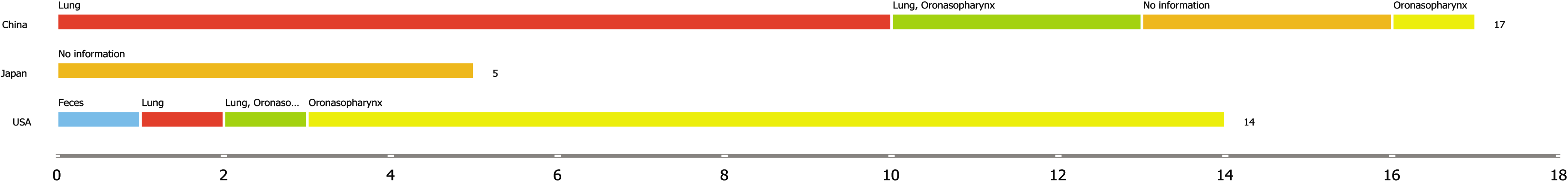
The distribution of samples across the sources of isolation (termed as isolates) and geographical location.

To further verify the significance of isolation sources, multiple classification algorithms were applied on the feature data with 5-fold cross validation. The same *n*-mer based normalized features (84 in total) were used. The cross validation results, in terms of classification accuracy, precision, recall, F1 score, and area under the ROC curve (AUC), are reported in Table 2. The classification results were not good based on the sources of isolation of the samples studied.

**Table 2:** Performance evaluation of different classification approaches (5-fold cross validation) based on the 84 *n*-mer features to distinguish the sources of isolation from where samples were collected. The best values over a column are shown in bold.

| Classification | Accuracy | Precision | Recall | F1 Score | AUC |
| --- | --- | --- | --- | --- | --- |
| k-NN | 41.7% | 0.409 | 0.417 | 0.407 | 0.669 |
| AdaBoost | 52.8% | 0.479 | 0.528 | 0.501 | 0.678 |
| Random Forest | **58.3%** | 0.502 | **0.583** | **0.539** | **0.800** |

On collecting the isolates of the samples, a significant overlap was found (Fig. 5). The observed frequencies of the geographical location and isolate in these samples indicate that the isolate is not an important feature to distinguish (*N* = 36, *χ*^2^ = 42.45, *p* = 0) the coronavirus samples. Hence, there is still no affinity towards the collection of samples from different body parts in human.

**Figure 5:**
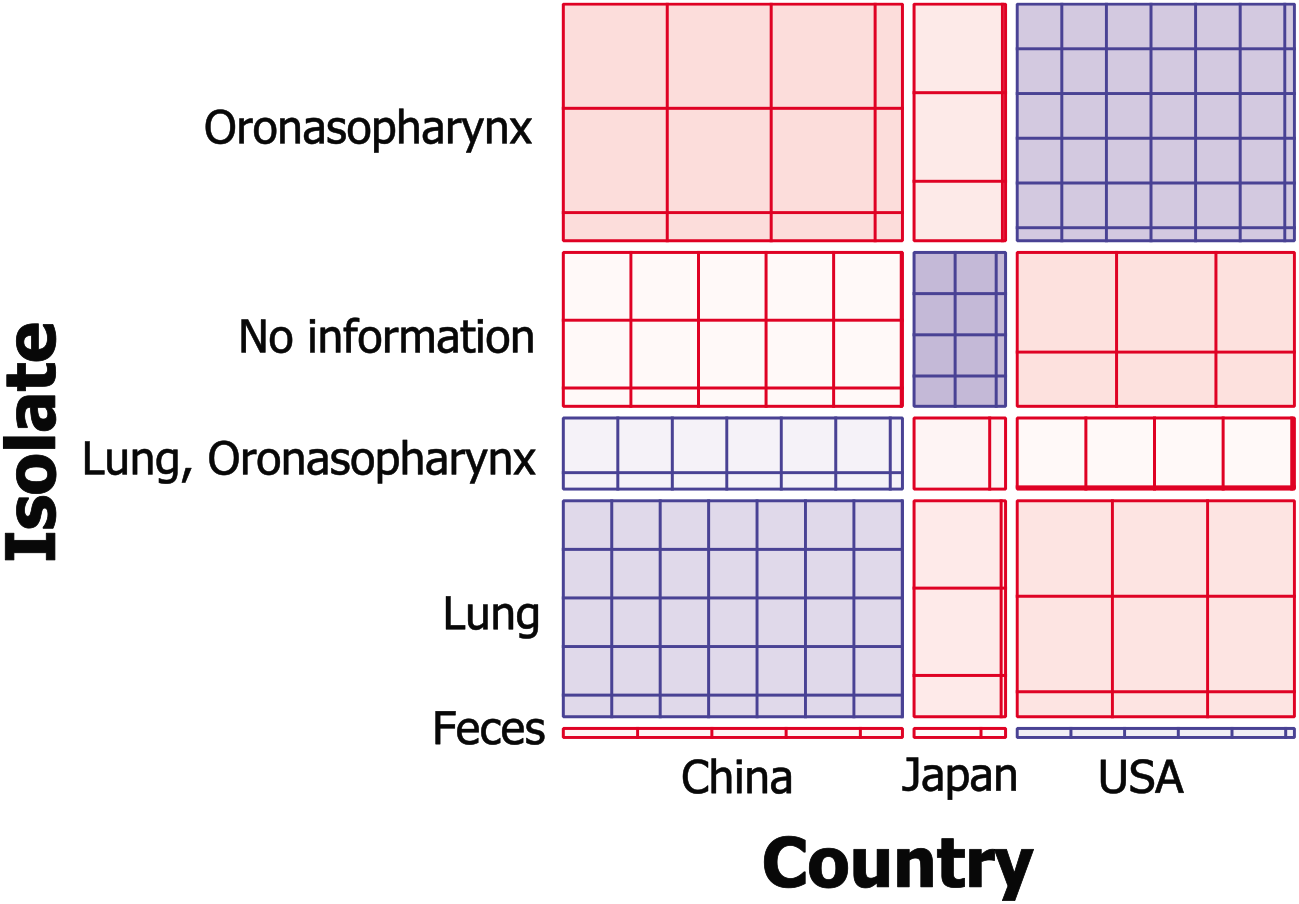
The Sieve plot showing distribution of samples in different geographical locations (marked as Country) versus different isolates (marked as Isolate).

### 4.4 Phylogenetic Analaysis

The phylogenetic tree was constructed based on the 36 sequences of the SARS-CoV-2 samples that we studied against the other samples of coronaviruses (Fig. 6). The results highlight a strong overlap of SARS-CoV-2 with human SARS and bat coronaviruses. Most interestingly, some of the sample sequences of SARS-CoV-2 collected from USA exhibit a separate phylogenetic closeness than the others. In fact, in a majority of the cases, the samples from the same location comes under the same branch in this tree, thereby highlighting a evolutionary connection between the sample and its collection footprint.

**Figure 6:**
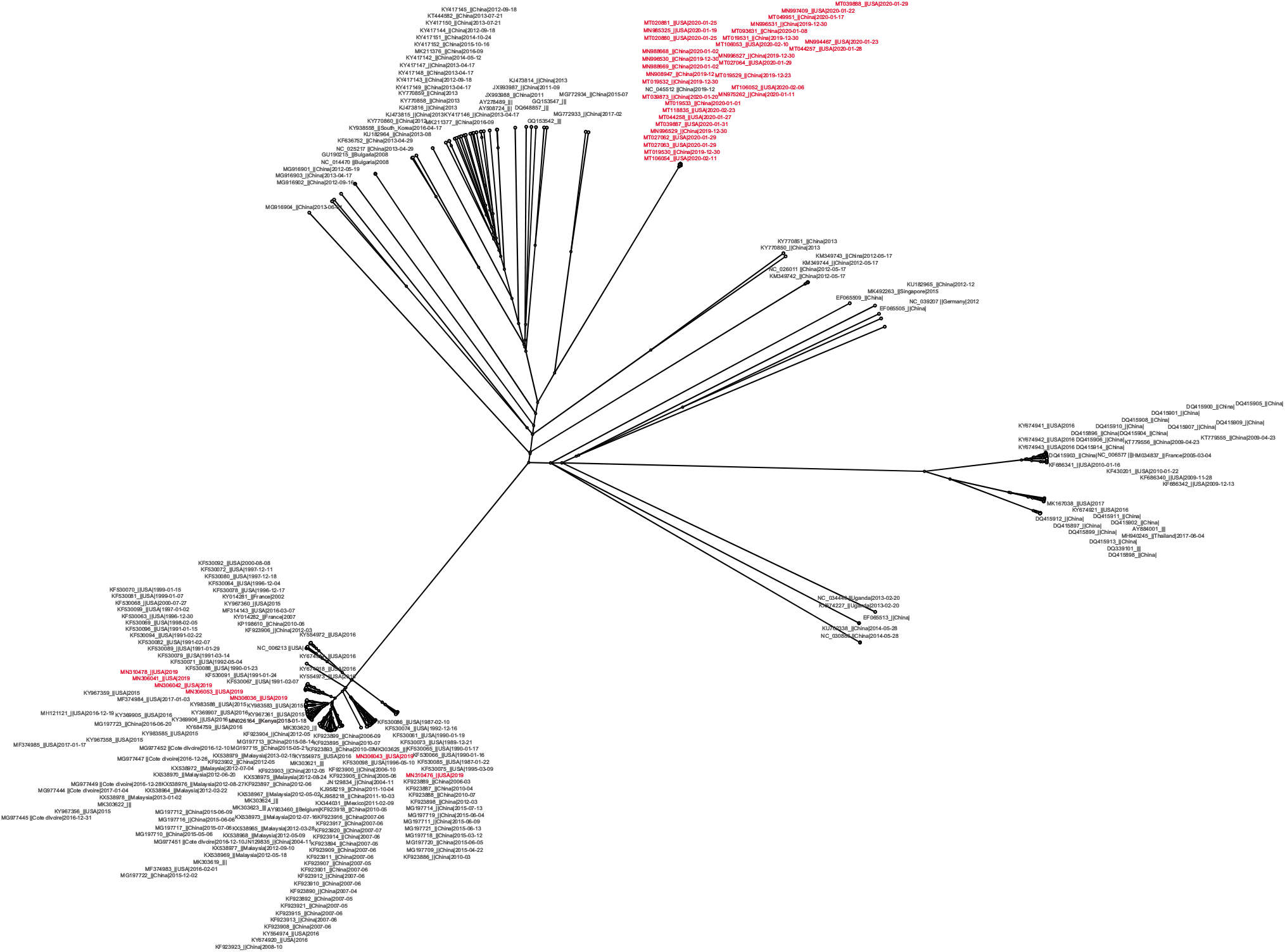
The phylogenetic tree connecting SARS-CoV-2 and other coronaviruses. The SARS-CoV-2 samples are marked in red. Some samples from USA form a group on the left side and the rest (a mixture of samples from USA and China) form a group on the right side.

### 4.5 Protein Analysis

On comparing the genomic sequence of Wuhan seafood market pneumonia virus isolate Wuhan-Hu-1 complete genome (29903 nt) with the protein sequence data bank UnitProt, two major proteins were found to match with many different coronaviruses. These include human, bat, middle east respiratory syndrome-related, murine, bovin, avian, porcine transmissible gastroenteritis, and feline coronaviruses, with bat coronaviruses having a major share (see Supplementary Fig. 1). Many of these are in fact experimentally validated (PE=1 as marked in UniProt). Interestingly, out of the three proteins found, two (Replicase polyprotein 1a and 1ab) were aligned to many coronaviruses but the third one (spike glycoprotein) was aligned to human SARS coronavirus only. These proteins are now being studied in many recent studies to link mutations in them with the pathophysiology of SARS-CoV-2 [17]. The fast family and domain prediction results (see Supplementary Fig. 2) also highlight a deep connection between human SARS coronavirus and SARS-CoV-2.

## 5 Conclusion

The SARS-CoV-2 is still a poorly characterized virus due to its diverse genomic alterations, rapid power of proliferation from host to host, and spread over several countries. Due to such complex characterization, understanding the pathophysiology of SARS-CoV-2 can be a real challenge. As the samples from China and USA had distinct isolates but they were closer together in terms of *n*-mer distribution, hence there are other important factors like temporal shift of locations where the spread took place. This highlights a high complexity of understanding the effect of mutations taking place in this virus over time. It is equally interesting to explore how much dependent such mutations are on the geographic origin of the samples. As the recurrent occurrence of this virus in human is already characterized by its diversity [15], its aberrant mutations in different locations raise the possibility of its outbreak in future.

## Abbreviations

SARS-CoV-2: 2019 novel coronavirus
SARS-CoV: Severe Acute Respiratory Syndrome
MERS-Cov: Middle East Respiratory Syndrome
COVID-19: 2019 novel coronavirus disease.

## Consent for publication

Not applicable.

## Ethics approval and consent to participate

Not applicable.

## Availability of data and materials

All datasets used in this paper are freely available from the NCBI data repository. The data with normalized *n*-mer features is provided as Supplementary Dataset 1. The metadata for all the 87 sequences available and for those 36 that were considered in this analysis are provided as Supplementary Dataset 2 and 3, respectively.

## Authors’ contributions

SoB, SS and MB designed the experiments. SoB and SS carried out the experiments and data analysis. SoB, SS, SaB and MB drafted the manuscript. All the authors read and approved the final manuscript.

## Acknowledgments

SoB acknowledges Digital India Corporation (formerly Media Lab Asia), Ministry Of Electronics and Information Technology (MeitY), Government of India, for providing him a Senior Research Fellowship under the Visvesvaraya Ph.D. scheme for Electronics and IT.

## Condolence

The authors feel deep grief and sympathy for those who got affected by the SARS-CoV-2.

## Footnotes

1 genes/genomes http://covdb.microbiology.hku.hk

2 https://www.ncbi.nlm.nih.gov/labs/virus (accessed in February, 2020)

3 https://www.ncbi.nlm.nih.gov/nuccore

4 https://orange.biolab.si

## References

[1] Anthony R Fehr and Stanley Perlman. Coronaviruses: an overview of their replication and pathogenesis. In Coronaviruses, pages 1–23. Springer, 2015.

[2] Susan R Weiss and Julian L Leibowitz. Coronavirus pathogenesis. In Advances in Virus Research, volume 81, pages 85–164. Elsevier, 2011.

[3] Shuo Su, Gary Wong, Weifeng Shi, Jun Liu, Alexander CK Lai, Jiyong Zhou, Wenjun Liu, Yuhai Bi, and George F Gao. Epidemiology, genetic recombination, and pathogenesis of coronaviruses. Trends in Microbiology, 24(6):490–502, 2016.

[4] Nicole R Sexton, Everett Clinton Smith, Hervé Blanc, Marco Vignuzzi, Olve B Peersen, and Mark R Denison. Homology-based identification of a mutation in the coronavirus rna-dependent rna polymerase that confers resistance to multiple mutagens. Journal of virology, 90(16):7415–7428, 2016.

[5] W Tan, X Zhao, X Ma, W Wang, P Niu, W Xu, GF Gao, and G Wu. A novel coronavirus genome identified in a cluster of pneumonia cases—wuhan, china 2019-2020. China CDC Weekly, 2(4):61–62, 2020.

[6] Na Zhu, Dingyu Zhang, Wenling Wang, Xingwang Li, Bo Yang, Jingdong Song, Xiang Zhao, Baoying Huang, Weifeng Shi, Roujian Lu, et al. A novel coronavirus from patients with pneumonia in china, 2019. New England Journal of Medicine, 2020.

[7] Sasmita Poudel Adhikari, Sha Meng, Yu-Ju Wu, Yu-Ping Mao, Rui-Xue Ye, Qing-Zhi Wang, Chang Sun, Sean Sylvia, Scott Rozelle, Hein Raat, et al. Epidemiology, causes, clinical manifestation and diagnosis, prevention and control of coronavirus disease (covid-19) during the early outbreak period: a scoping review. Infectious diseases of poverty, 9(1):1–12, 2020.

[8] World Health Organization et al. Novel coronavirus (2019-ncov): situation report, 94. 2020.

[9] Eleanor M Cottam, Jemma Wadsworth, Nick J Knowles, and Donald P King. Full sequencing of viral genomes: practical strategies used for the amplification and characterization of foot-and-mouth disease virus. In Molecular Epidemiology of Microorganisms, pages 217–230. 2009.

[10] Deepak Sharma, Pragya Priyadarshini, and Sudhanshu Vrati. Unraveling the web of viroinformatics: computational tools and databases in virus research. Journal of Virology, 89(3):1489–1501, 2015.

[11] Kai Kupferschmidt. Preprints bring ‘firehose’ of outbreak data. Science, 367(6481):963–964, 2020.

[12] Roujian Lu, Xiang Zhao, Juan Li, Peihua Niu, Bo Yang, Honglong Wu, Wenling Wang, Hao Song, Baoying Huang, Na Zhu, et al. Genomic characterisation and epidemiology of 2019 novel coronavirus: implications for virus origins and receptor binding. The Lancet, 395(10224):565–574, 2020.

[13] Carmine Ceraolo and Federico M Giorgi. Genomic variance of the 2019-ncov coronavirus. Journal of Medical Virology, 2020.

[14] Jiahua He, Huanyu Tao, Yumeng Yan, Sheng-You Huang, and Yi Xiao. Molecular mechanism of evolution and human infection with sars-cov-2. Viruses, 12(4):428, 2020.

[15] Jie Cui, Fang Li, and Zheng-Li Shi. Origin and evolution of pathogenic coronaviruses. Nature Reviews Microbiology, 17(3):181–192, 2019.

[16] Yi Huang, Susanna KP Lau, Patrick CY Woo, and Kwok-yung Yuen. CoVDB: a comprehensive database for comparative analysis of coronavirus genes and genomes. Nucleic Acids Research, 36(suppl_1):D504–D511, 2007.

[17] Aiping Wu, Yousong Peng, Baoying Huang, Xiao Ding, Xianyue Wang, Peihua Niu, Jing Meng, Zhaozhong Zhu, Zheng Zhang, Jiangyuan Wang, et al. Genome composition and divergence of the novel coronavirus (2019-ncov) originating in china. Cell Host & Microbe, 2020.

[18] Janez Demšar, Tomaž Curk, Ales Erjavec, Crt Gorup, Tomaž Hocevar, Mitar Miluti-novic, Martin Možina, Matija Polajnar, Marko Toplak, Anže Staric, et al. Orange: data mining toolbox in python. The Journal of Machine Learning Research, 14(1):2349–2353, 2013.

[19] Richard O Duda, Peter E Hart, and David G Stork. Pattern Classification. John Wiley & Sons, 2012.

[20] William R Pearson, Todd Wood, Zheng Zhang, and Webb Miller. Comparison of dna sequences with protein sequences. Genomics, 46(1):24–36, 1997.

[21] Barbara Holland and Vincent Moulton. Consensus networks: A method for visualising incompatibilities in collections of trees. In International Workshop on Algorithms in Bioinformatics, pages 165–176. Springer, 2003.

