## Supplementary figures and images for "Tracing Back the Temporal Change of SARS-CoV-2 with Genomic Signatures"

### Supplementary Fig. 1

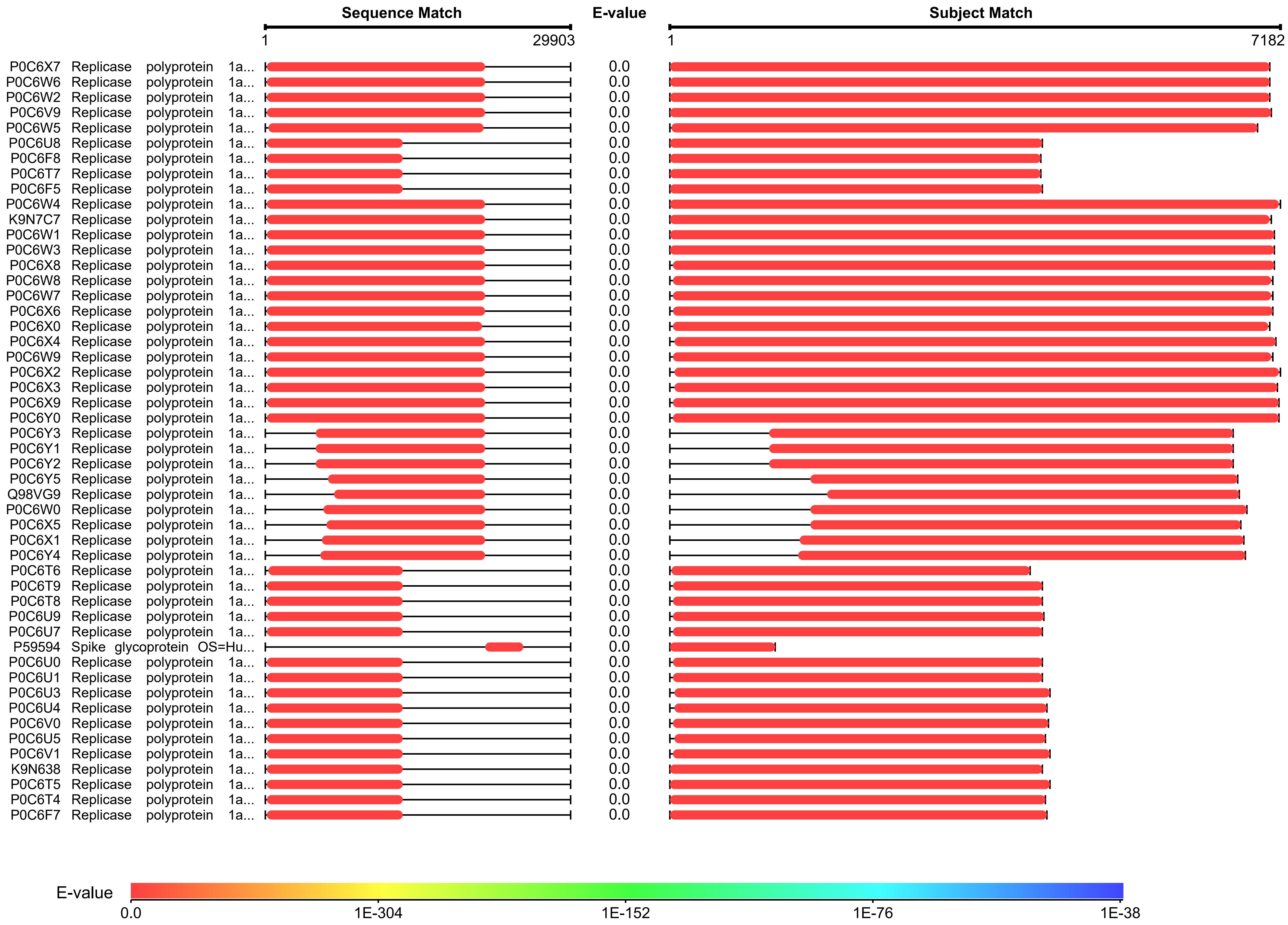

### Supplementary Fig. 2

**Fast Family and Domain Prediction**

(Query Sequence View)

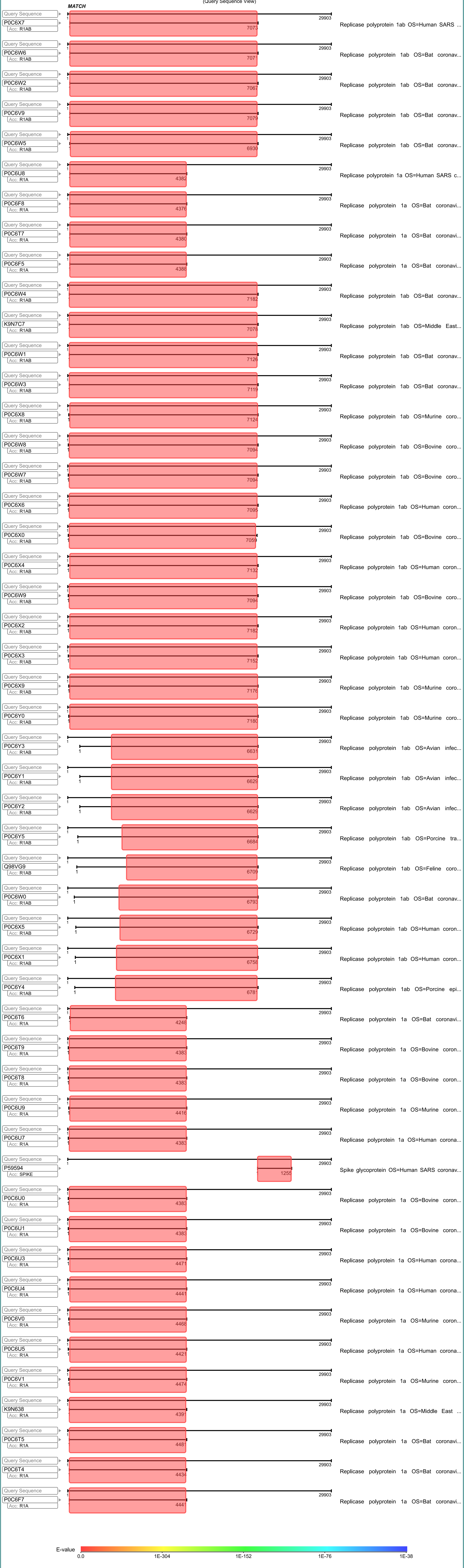
